# Genomic evaluation of BC4, a consortium of four *Alkalihalobacillus clausii* isolates, confirms its probiotic potential and safety in usage

**DOI:** 10.1101/2022.07.27.501716

**Authors:** Tanisha Dhakephalkar, Shilpa Wagh, Kunal Yadav, Anupama S. Engineer, Soham D. Pore, Prashant K. Dhakephalkar

## Abstract

Four strains of *Alkalihalobacillus clausii* B603/Nb (resistant to rifampicin), B619/R (resistant to streptomycin), B637/Nm (resistant to tetracyclin) and B106 (resistant to chloramphenicol) were isolated from various sources and used to prepare a consortium designated as BC4. Genomes of the constituent strains of the BC4 consortium were evaluated to investigate their genetic makeup and determine their probiotic potential. Gene prediction and functional annotation were performed using RAST. The data obtained was mined for genes encoding various phenotypic traits. This analysis revealed the presence of several genes encoding probiotic attributes like (i) survivability in the presence of low pH, bile, oxidative stress; (ii) bacterial aggregation and adhesion to gut epithelium, etc.; and (iii) enzymes/ molecules conferring health benefits. Further, the genome analysis also confirmed the genes required for enhancing the nutritional amenability, health-promoting, and disease-preventing traits were present. Several genes encoding multiple antibiotic resistance were detected; however, none of these genes was located on mobile elements such as plasmids, transposons, etc. The absence of genes encoding virulence factors, pathogenic islands, emetic toxins, etc., as well as mobile genetic elements, underscored the safety of BC4 isolates.

## INTRODUCTION

Probiotics are “live microorganisms which when administered in adequate amounts confer a health effect on the host” (FAO/WHO 2001). The use of Probiotics is mainly for the restoration of healthy gut microbiota. Often, during antibiotic treatment, the normal microbial flora associated with the gut gets disrupted. Hence, probiotics and oral antibiotics are prescribed together in clinical medicine to restore healthy gut microbiota (Kechagia et al., 2013). The most commonly used probiotics include members of the genus, *Lactobacillus, Saccharomyces*, and *Bifidobacterium*. However, most of these genera members cannot survive the gastric transit characterized by low pH, bile salts, presence of digestive juices such as amylase, trypsin, pepsin, etc. Thus, the potential probiotic strains must possess the mechanism to overcome such harsh physiological stress/ conditions.

Interestingly spore-forming bacteria have been increasingly gaining importance as probiotics over the last few decades. Endospore forming ability is associated with some aerobic and anaerobic rods and a few cocci. Endospores are metabolically dormant. They can resist relatively high or freezing temperatures, desiccation, toxic chemicals, and radiation. The ability of endospores to resist eco-physiological stress is an essential attribute for application in the probiotic industry.

Enterogermina (Sanofi Winthrop, Milan, Italy) has been an antidiarrheal probiotic in use for the last few decades (Mazza 1994). It comprises four strains of endospore-forming *Alkalihalobacillus clausii* that are resistant to multiple antibiotics (Ciffo, 1984). Interestingly, there is a lack of information on the probiotic characterization of four individual Enterogermina strains other than 16S rRNA gene sequence-based identification, antibiotic resistance determination, and immunomodulation. Enterogermina is not explicitly referred to as a probiotic but claims to enhance the body’s immune system following the germination of spores in the small intestine. Recently, the composite genome of Enterogermina was sequenced and characterized to gain insights into its probiotic potential (Khatri et al., 2019). However, the genome sequence of individual strains was not available in the public domain as per the WHO/FAO guidelines.

FAO/ WHO has recommended the following guidelines as minimal safety measures for probiotics composed of spore-forming microorganisms (SCAN 2000a, b; FAO/WHO 2001). Each bacterium in the product must be isolated and unequivocally identified using the most current, valid methodology. The strain nomenclature must be consistent with the present, scientifically recognized names. Each bacterial strain must be sufficiently characterized in vitro. Such characterization must include (i) profiling of antibiotic resistance, (ii) production of emetic or enterotoxins, and (iii) resistance to gastric acid and bile. One must not use bacterial strains possessing transferable antibiotic resistance. Probiotic formulations must not include strains capable of producing toxins.

The present investigation was undertaken to (i) isolate, identify and characterize the constituent strains of the BC4 consortium based on the differences in their antibiotic resistance profile; (ii) determine the ability of individual strains to survive gastric transit; (iii) sequence the whole genome of each strain and gain genomic insights into the probiotic potential of each of the isolates; (iv) investigate safety aspects of individual strains.

## Materials and Methods

### Bacterial strain, DNA preparation, and Genome Sequencing

*Alkalihalobacillus clausii* strains constituting the BC4 consortium were procured from Hi Tech BioSciences India Pvt. Ltd., Pune, India. Four isolates B603/Nb, B619/R, B637/Nm (this study), and B106, are mixed to form consortium BC4 (Kapse et al., 2019). All strains are deposited in National Center for Microbial Resource (NCMR), Pune, India, viz. MCC 0190, MCC 0189, MCC 0191 and MCC 0188. The genomes of B603/Nb, B619/R, and B637/Nm, were sequenced using the Illumina HiSeq PE platform consisting of four basic steps, viz. sample preparation, library construction, sequencing, and data generation. Raw data was generated using an integrated primary analysis software called RTA (Real-Time Analysis). The type of reading was paired-end, and the read length was 151 bp. Sequencing was outsourced to Macrogen, South Korea. The raw data obtained was in the form of FASTQ files which were used for genome assembly and further data analysis.

### Genome assembly and annotation

De novo genome assembly was carried out using SPAdes assembler version 3.9.1 (Bankevich et al., 2012). Functional annotation of the B603/Nb, B619/R, B637/Nm, and B106 genomes was carried out by the Rapid Annotation using Subsystem Technology, abbreviated as RAST (Aziz et al., 2008). The genome was mined for the presence of probiotic marker genes associated with health promotion and disease prevention using the RAST server. The individual probiotic attributes are described along with their FIGfam numbers. FIGfam is a set of proteins that are believed to be isofunctional homologs, having the same function, and are derived from a common ancestor.

### Comparative genome analysis

The whole-genome shotgun project of the *Alkalihalobacillus clausii* B603/Nb, B619/R, B637/Nm, and B106 has been deposited at DDBJ/ENA/GenBank under the accession JABFCU000000000, JABFCW000000000, JABFCV000000000 & NFZO00000000 respectively. The genome sequence of ENTpro (composite genome of Enterogermina formulation) was obtained from the NCBI database for the comparative genome analysis of the *A. clausii* B603/Nb, B619/R, B637/Nm, and B106. Digital DNA DNA hybridization for *in-silico* genome to genome comparison was carried out as described by Auch et al. (2010), using the genomes of B603/Nb, B619/R, B637/Nm, and B106 as a reference, with the help of online tool http://ggdc.dsmz.de/. Blast Ring Image Generator (BRIG) software was used to generate a comparative circular genome image. The genome sequences of query and reference were submitted in the ’fasta’ format to generate the image with ENTpro as the reference strain. Nucleotide alignments of multiple query genomes against the reference strain were generated using BLASTn to create the circular genome comparison map (Alikhan et al., 2011).

### Prediction of secondary metabolite and bacteriocin production

antiSMASH webserver was used to predict the antibiotic and secondary metabolite biosynthetic gene clusters from the genome of B603/Nb, B619/R, B637/Nm, and B106. antiSMASH uses a large number of in silico secondary metabolite analysis tools, integrates them, and provides the prediction (Weber et al., 2015). The bacteriocin mining was executed in BAGEL 3 web-based server using the ’fna’ file from the four *A. clausii* strains. The software works with input data evaluated against a curated dataset of bacteriocins (van Heel et al., 2013).

### Antibiotic resistance prediction

Antibiotic resistance genes in the genomes of four *A. clausii* isolates were predicted using the web-based server CARD, a thoroughly curated collection of identified resistance genes and related antibiotics by the Antibiotic Resistance Ontology (ARO) and AMR gene detection models. CARD analyses the genome sequences using BLAST and the Resistance Gene Identifier (RGI) software for prediction (Jia et al., 2017).

### Results and Discussion

It is of paramount importance to unequivocally establish the identity of each probiotic strain up to the species level. Hence, in the present investigation, four strains constituting the BC4 consortium were characterized based on antibiotic resistance and identified using advanced molecular tools.

### Identification of BC4 strains

Four BC4 consortium strains, designated as B603/Nb (resistant to Novobiocin), B619/R (resistant to rifampicin), B637/Nm (resistant to Neomycin) (this study), and B106 (resistant to tetracyclin) (Kapse et al., 2019), that differed in terms of their antibiotic resistance profile were isolated by and procured from Hi Tech BioSciences India Pvt. Ltd.. Each of these four isolates formed large, creamy white, opaque colonies with irregular margins on Nutrient Agar medium on incubation for 72 hrs at 37°C. All four isolates were Gram-positive, spore-forming, and non-motile long rods, which often occurred in chains and/or clumps. Biochemical identification/ characterization of B603/Nb, B619/R, B637/Nm, and B106 was carried out using the Biolog’s advanced phenotypic technology. All four isolates displayed comparable biochemical profiles and were identified as members of *Alkalihalobacillus clausii*. The identity of each isolate was further confirmed by sequencing the 16S rRNA gene. Each of the four strains shared more than 99.85% 16S rRNA gene sequence homology with the closest phylogenetic affiliate, *Alkalihalobacillus clausii* DSM 8716. Each of the four isolates shared 99.85% 16S rRNA gene sequence homology with four reference sequences of Enterogermina strains (O/C, SIN, N/R, and T) available in the GenBank database. The whole-genome shotgun project of the four isolates, namely *Alkalihalobacillus clausii* B603/Nb, B619/R, B637/Nm, and B106, has been deposited at DDBJ/ENA/GenBank under the accession JABFCU000000000, JABFCW000000000, JABFCV000000000 (This study) & NFZO00000000 (Kapse et al., 2019) respectively. Identification of BC4 consortium strains was further confirmed with the help of digital DNA-DNA hybridization (DDH) to determine the genome homology of BC4 consortium strains with reference and other probiotic *A. clausii* genome sequences available in the GenBank database (Table 1). All four BC4 strains shared almost 100% DDH values. Interestingly, DNA DNA Hybridization of the three isolates included in this study with type strain *A. clausii* DSM 8716 was only 62.50% when calculated with the recommended formula (Auch et al., 2010). However, these isolates shared more than 95% DDH value with probiotic *A. clausii* B106, ENTPro, UBBC-07.

**Table 1:** DNA homology between BC4 strains and reference/ probiotic *A. clausii* genomes determined by Digital DNA-DNA hybridization

|  | **B603Nb** | **B619R** | **B637Nm** | **B106** |
| --- | --- | --- | --- | --- |
| **UBBC-07** | 92.00% | 92.00% | 92.00% | 100% |
| **KSM-K16** | 94.30% | 94.30% | 94.30% | 100% |
| **ENTPro** | 99.30% | 99.20% | 99.30% | 100% |
| **AKU0647** | 50.70% | 50.70% | 50.70% | 50.60% |
| **DSM8716** | 62.50% | 62.50% | 62.50% | 62.50% |

To establish the unequivocal identity of BC4 strains, the multigene sequence homology of these isolates with the reference sequences in the GeneBank database was determined. It was observed that the BC4 strains shared the highest homology for each of the housekeeping gene sequences with the corresponding genes of reference, *A. clausii* DSM 8716 (Table 2). The sequence homology for each gene between BC4 strains and *A. clausii* was more than 96%. These observations underscored the unequivocal affiliation of BC4 strains with *A. clausii* species. It was noteworthy that BC4 strains shared a genome sequence homology of more than 99% with ENTPro, the composite genome of Enterogermina reported earlier in an unrelated study (Khatri et al., 2010).

**Table 2:**
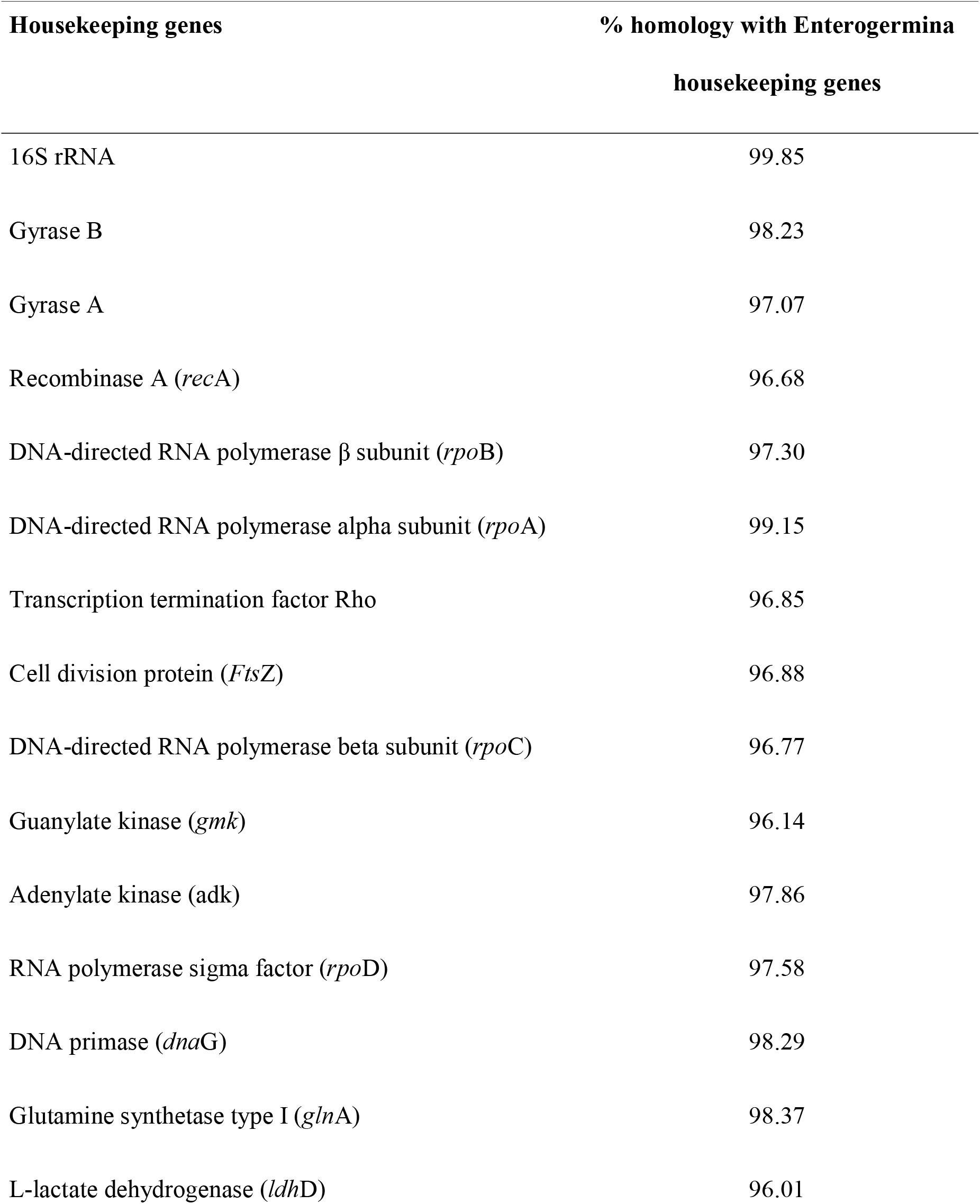
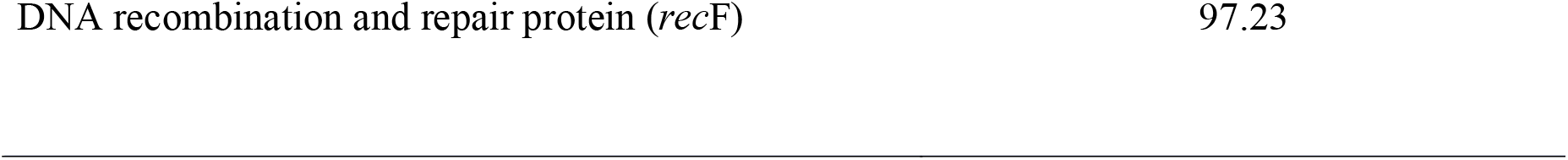
Homology between BC4 strains (this study) and *A. clausii* DSM8716 housekeeping genes.

### Genomic features of *Alkalihalobacillus clausii* strains B603Nb, B619R, B637Nm, and B106

Illumina-based sequencing reads were assembled using SPAdes assembler version 3.9.1 (Bankevich et al., 2012). The resultant contigs for each isolate are illustrated in (Table 3). Contigs for each strain were assembled to form a presumptive circular chromosome of approximately 4.24 Mb, a size that was comparable for all four stains. The genome of strain B603Nb was 4,244,642 bp from 22 contigs, the genome of strain B619R was 4,243,844 bp from 21 contigs, the genome of strain B637Nm was 4,243,991 bp from 21 contigs. The GC content for all three strains was observed to be 44.7% and had an N50 of 629869. N50 refers to the minimum contig length needed to cover 50% of the genome. A total of 317 subsystems, 4466 protein-coding sequences, and 91 RNA genes were predicted using RAST (Aziz et al., 2008). Out of the total RNA genes, 75 were found to be of tRNA, and 16 were of rRNA after annotating the genome using PATRIC, as described by Brettin et al. (2015). The circular image comparing the genomes of probiotic-and reference *A. clausii* strains were constructed using the online tool BRIG (BLAST Ring Image Generator). The genome of B603/Nb was used as a reference. Close to 100% homology was observed amongst the genomes of the four BC4 strains (Figure 1). Similarly, the composite ENTPro genome also shared 99.30% genome homology with the BC4 strains obtained in the present study. Some differences were observed between genome sequences of UBBC-07 and DSM-8716. Negligible gaps and high homology indicated the completeness of the genome sequences of the four BC4 strains.

**Table 3:** Genome features of assembled genomes of BC4 strains (This study)

| Genome features |  |  |  |
| --- | --- | --- | --- |
|  | B603Nb | B619R | B637Nm |
| Genome size (bp) | 4,244,642 | 4,243,844 | 4,243,991 |
| GC content (%) | 44.7 | 44.7 | 44.7 |
| Protein coding sequences (CDS) | 4466 | 4468 | 4466 |
| Subsystems | 317 | 317 | 317 |
| tRNA | 75 | 75 | 75 |
| rRNA | 16 | 16 | 16 |
| Number of contigs | 22 | 21 | 21 |
| Hypothetical proteins | 1536 |  |  |

**Figure 1.**
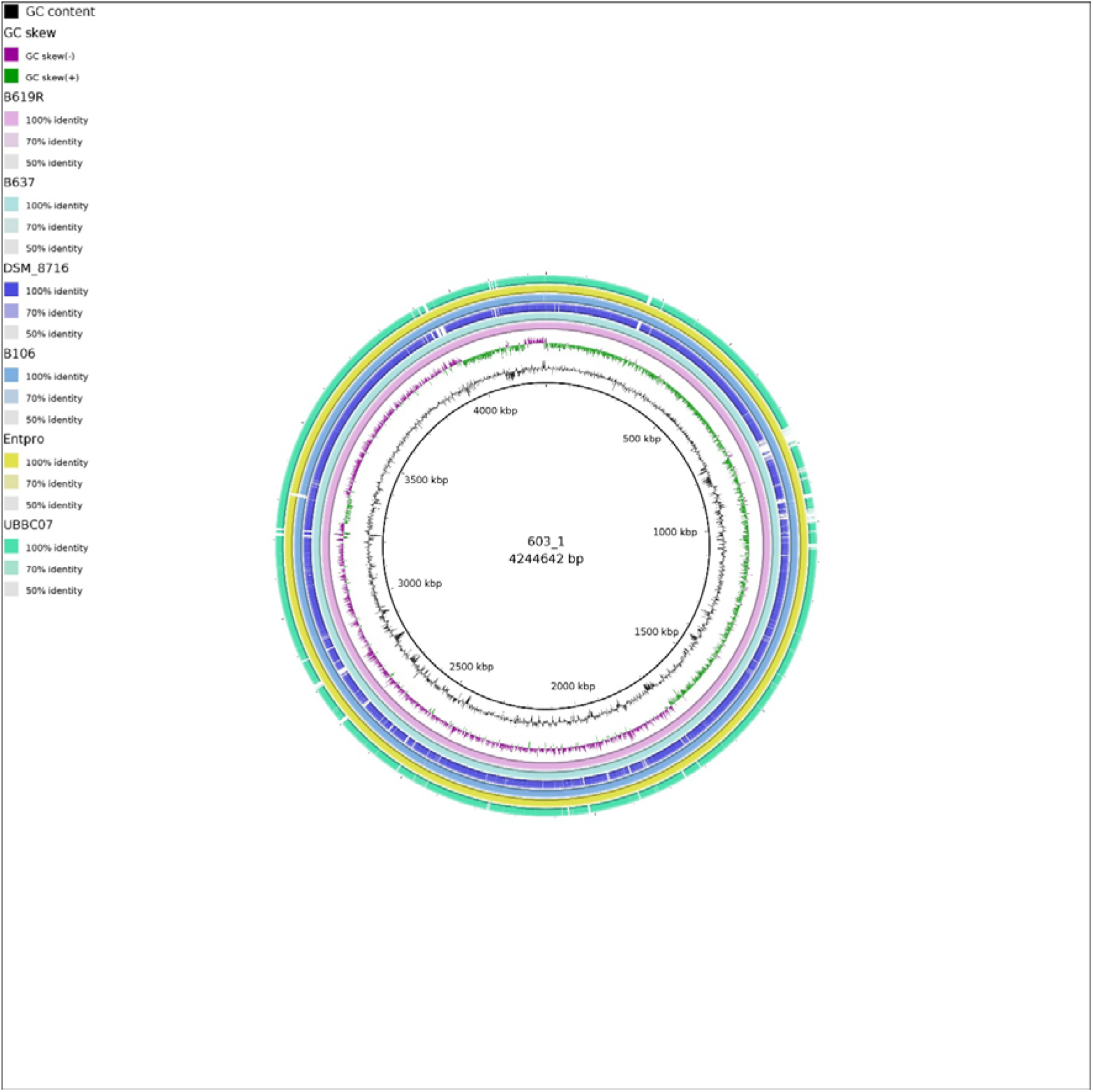
Circular comparison of genomes of probiotic and reference *A. clausii* isolates with the B603Nb genome as a reference. Each colored ring represents a query genome. The intensity of the color represents relative levels of nucleotide homology between the reference and query genomes.

**Figure 2:**
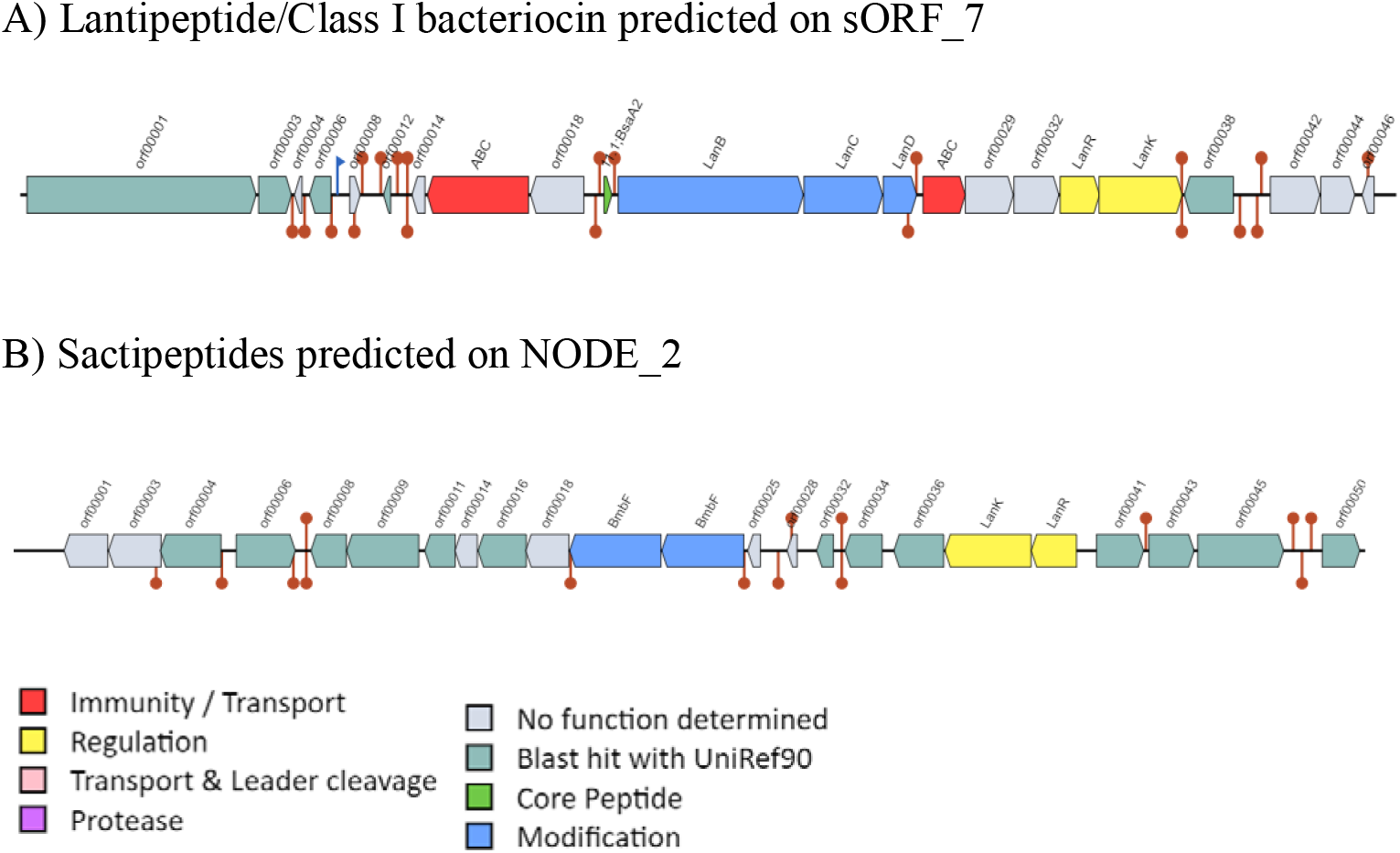
Genetic organization of putative bacteriocins from the genomes of *A. clausii* strains B603Nb, B619R, B637Nm, and B106

### Antibiotic resistance profile of the *A. clausii* strains B603/Nb, B619/R, B637/Nm, and B106

Probiotics, which are resistant to multiple antibiotics, can be co-administered with antibiotics to restore the healthy microbiota in the human gut. Such an oral probiotic supplement can prevent antibiotic-induced diarrhea in patients under antibiotic treatment. The present experimental data provided an updated antibiotic susceptibility profile of the four BC4 strains (Table 4). Each of the four strains was resistant to multiple antibiotics belonging to penicillins, cephalosporins, fluoroquinolones, aminoglycosides, and macrolides. Antibiotic resistance of the BC4 strains was similar to the Enterogermina isolates as described in the past (Ciffo, 1984; Mazza, 1992; Abbrescia et al., 2014). The resistance of BC4 strains observed in the present study against multiple antibiotics belonging to cephalosporins, quinolones, macrolides, aminoglycosides, and penicillins was consistent with the antibiotic resistance profile of the Enterogermina strains reported earlier by Abbrescia et al. (2014).

**Table 4:**
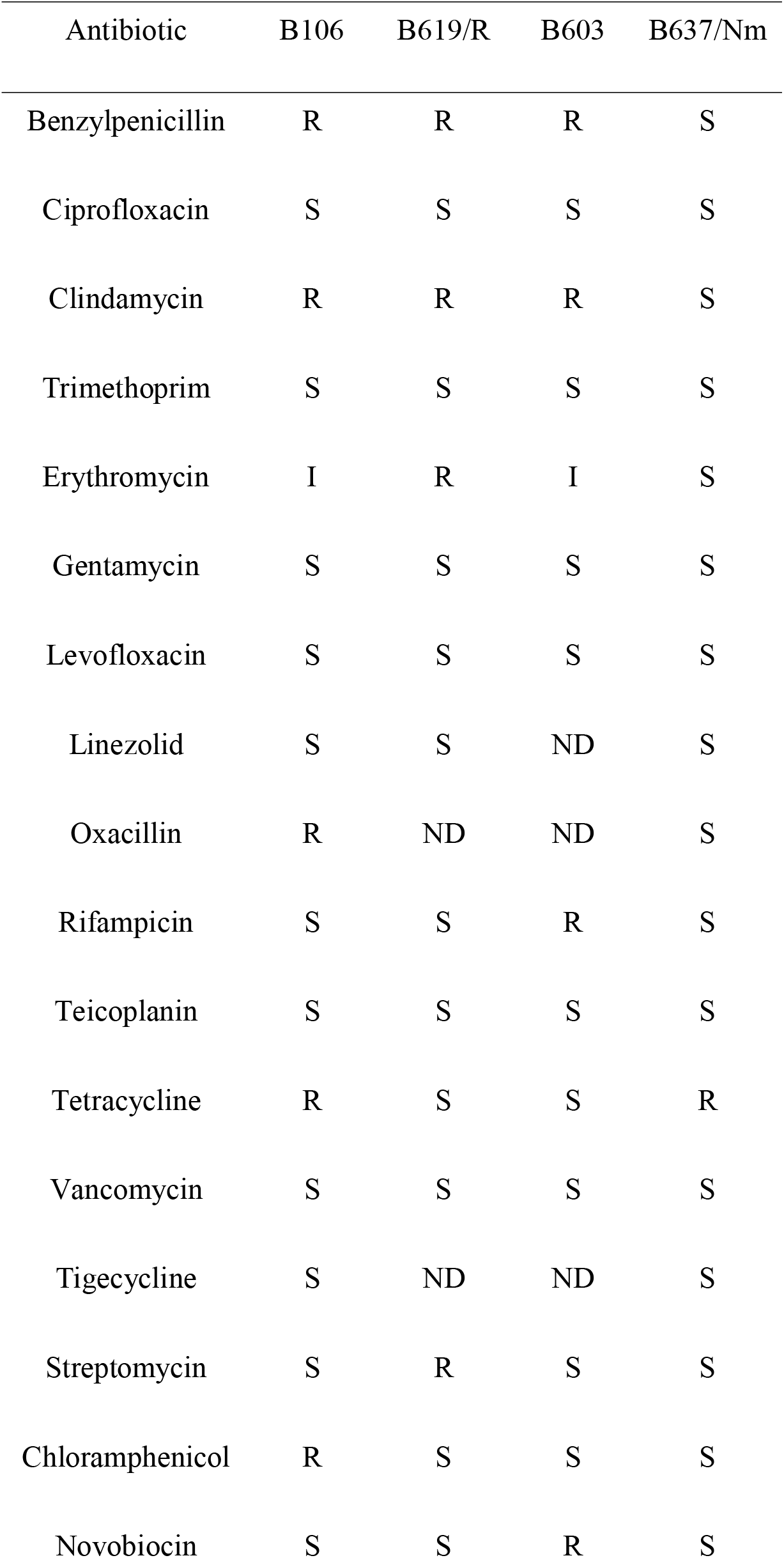
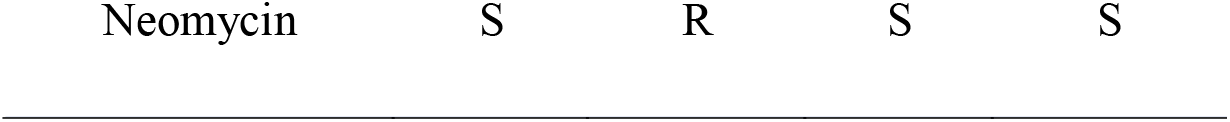
Resistance of BC4 strains to antibiotics.

In the present investigation, attempts were made to gain insight into the genetics of antibiotic resistance mechanisms and to determine the intrinsic nature or otherwise of the genes involved. Bacteria can become resistant to antibiotics by two mechanisms: i) occurrence of mutations in essential (housekeeping) genes. In such cases, the resistance is intrinsic and not transferable. ii) procurement of an external resistant determinant from another bacterium via horizontal gene transfer. This type of acquired resistance is re-transferable. The resistance of *A. clausii* strains to multiple antibiotics enables them to survive in the presence of the antibiotics (Courvalin, 2006; Abbrescia, et al., 2014). The present investigation provided an overview of the antibiotic susceptibility pattern amongst the four BC4 strains. Genetic analysis of these isolates revealed various genetic factors that could contribute to antibiotic resistance. Mining of the genomes of four isolates, B603/Nb, B619/R, B637/Nm, and B106, revealed the presence of genes encoding for classes A, C, and D beta-lactamases and aminoglycoside nucleotidyltransferase. Chromosomal resistance to tetracyclines could be attributed to the presence of ’ribosome protection’ type tetracycline resistance protein which was detected in the genome of B637/Nm. Ribosome protection proteins are members of the translation factor superfamily of GTPases that bring about the release of tetracycline from the ribosome in a GTP-dependent manner which decreases drug binding (Wilson, 2014). Resistance to chloramphenicol could be ascribed to the genes encoding for chloramphenicol acetyltransferase found in the genome of strain B106 (Kapse et al., 2019). This enzyme catalyzes the transfer of an acetyl group from acetyl Co A to the chloramphenicol molecule, thereby preventing chloramphenicol from binding to the ribosome (Cm et al., 1982). Abbrescia et al., 2014 reported resistance to rifampicin was due to a chromosomal mutation (a change in amino acid sequences resulting in the S488F missense) in the rpoB gene encoding the β subunit of RNA polymerase. A similar observation was obtained in the genome of B603/Nb (a change in amino acid sequences resulting in the S491F missense).

. One potential mechanism which can be responsible for the resistance to streptomycin is as follows: Aminoglycoside nucleotidyltransferases (ANTs) mediate the inactivation of aminoglycosides by catalyzing the transfer of an AMP group from the donor substrate ATP to a hydroxyl group in the aminoglycoside molecule. Out of the five classes of ANTs that catalyze adenylation, two classes were detected in the genome of B619/R, namely Aminoglycoside 6-nucleotidyltransferase and Aminoglycoside 4’-nucleotidyltransferase. This provided genomic evidence which correlated with the phenotypic results observed in the case of B619/R. The BC4 strains showed resistance to erythromycin as well. This property is probably due to the presence of the gene *erm34*, which encodes for macrolide resistance. The *erm* gene codes for a methylase which imparts resistance through ribosome methylation. The chromosomal location of this gene was indicative of its non-transferable nature, as was reported in the previously published study, which reported that the attempts to transfer the *erm* gene by conjugation were unsuccessful (Bozdogan et al., 2004).

### Evaluation of safety in the usage of BC4 strains

**P**robiotics must be safe for consumption; hence, certain concerns need to be addressed before any organism or strain is selected as a probiotic. Some of the main concerns revolving around the safety aspects of a probiotic are: i) they shouldn’t contain any plasmids or transposable elements which might aid in the transfer of antibiotic resistance determinants to the gastrointestinal flora, ii) they shouldn’t produce any toxin, and iii) the strain in question should not cause any diseases (Sharma et al., 2014). It is important to point out that many of the genetic determinants encoding antibiotic resistance, virulence factors, etc., are often located on mobile elements, such as transposons and plasmids. Analysis of genomes of BC4 strains with the help of Plasmid Finder Tools did not reveal the presence of any plasmids associated with the genome sequences of any of the four isolates investigated in the present study.

The genomes of BC4 strains were also assessed for the presence of pathogenicity islands and virulence factors using the Virulence Finder 2.0 server (Joensen et al., 2014). Genes or their variants encoding virulence factors in *S. aureus, Enterococcus, Listeria*, and *Escherichia coli* were absent in genomes of all four BC4 strains. The genomes of *A. clausii* strains B603/Nb, B619/R, B637/Nm, and B106 were also mined to check the presence of genes encoding toxins using the sequences of emetic toxin genes downloaded from the GenBank database as references. Such analysis confirmed the absence of the following genes in the genomes of any of the four BC4 strains: (i) bceT gene, encoding the single-component enterotoxin T; (ii) hblA, hblB, and hblC genes, which encode the hemolytic enterotoxin hbl; (iii) nheA, nheB and nheC genes which encode non-hemolytic enterotoxin (Nhe). Thus, the absence of any (i) mobile genetic elements, (ii) genes encoding virulence factors or emetic toxins, and (iii) pathogenic islands underscored the safety of BC4 strains investigated in the present study as probiotics. To our best knowledge, this is the first report of genetic evaluation of the safety of potential probiotics *A. clausii* strains that are genetically similar to Enterogermina isolates.

### Comparative analysis of genomes of probiotic and reference *A. clausii* strains

It was desired in the present study to characterize the genomes of each of the four BC4 strains and evaluate any minor differences they may have in their genetic makeup. Gene prediction and functional annotation were performed using RAST. The data obtained was mined for genes encoding various phenotypic traits. This analysis was undertaken to investigate the presence of several genes encoding (i) probiotic attributes like survivability in the presence of low pH, bile, and oxidative stress; (ii) bacterial adhesion and aggregation; (iii) enzymes that enhance the nutritional amenability; (iv) health-promoting and disease-preventing traits, etc. (V)industrially important enzymes/ molecules.

### Evaluating the potential of BC4 strains to survive low pH

Coping with acidic environments is one of the prerequisites for any probiotic organism. Several factors are involved in the adaptation of microorganisms to such acidic environments. One such factor is the presence of sigma B, which is an alternative sigma factor. Alternative sigma factors along with core RNA polymerases can alter global gene expression. Sigma B contributes to resistance to the acidic environment by two mechanisms: firstly, a general acid tolerance to which sigma B-regulated systems contribute throughout all growth phases, and secondly via a pH-inducible ATR mechanism that is partially regulated by sigma B-in exponential phase cells (Ryan et al., 2008). Thus, the presence of this factor is significant in the mechanism of acid tolerance in probiotic bacteria. Another factor that helps bacteria survive in acidic conditions is the ADI (arginine deaminase pathway). In a classical pathway, bacteria can utilize the extracellular arginine from the host’s diet. This arginine is received by the ADI system through the help of arginine/ornithine antiporter (Cotter & Hill 2003). Another interesting observation to note is that the ADI pathway is under the control of sigma factor B (Hain et al., 2008). Sigma B factor, arginine/ornithine antiporter ArcD, Ornithine carbamoyl-transferase, and Arginine pathway regulatory protein ArgR, a repressor of arg regulon, were all found to be present in all four BC4 strains. Expression of chaperones, such as GroES, GroEL, Dnak, DnaJ, and clp proteases which are known to be involved in protein repair mechanisms, were detected. Other transcriptional regulators like CstR, HrcA, and Crp, which are known to change upon exposure to low pH, were present. Some genes which code for symporters that allow bacteria to thrive in the acidic conditions of the gastrointestinal tract were also found. Some of these symporters include Na+/H+ antiporter (4 copies), H+/glutamate symporter, Glucose/mannose: H+ symporter, and Amino acid/H+ symporter (Lehri et al., 2017). On exposure to low pH, the electron transfer chain (ETC) is severely disturbed, which leads to the formation of reactive oxygen species (ROS). To counteract these, ROS inducement of oxidative stress mechanisms (Schieber & Chandel, 2014) may take place, including thioredoxins (proteins that act as antioxidants), catalase, superoxide dismutase, etc., all of which are present in all four strains (Mols & Abee, 2011). The presence of the genes summarized in Table 5 indicates the ability of the strains to survive the harsh conditions encountered in the GI tracts, like acidic pH and the presence of bile salts.

**Table 5:**
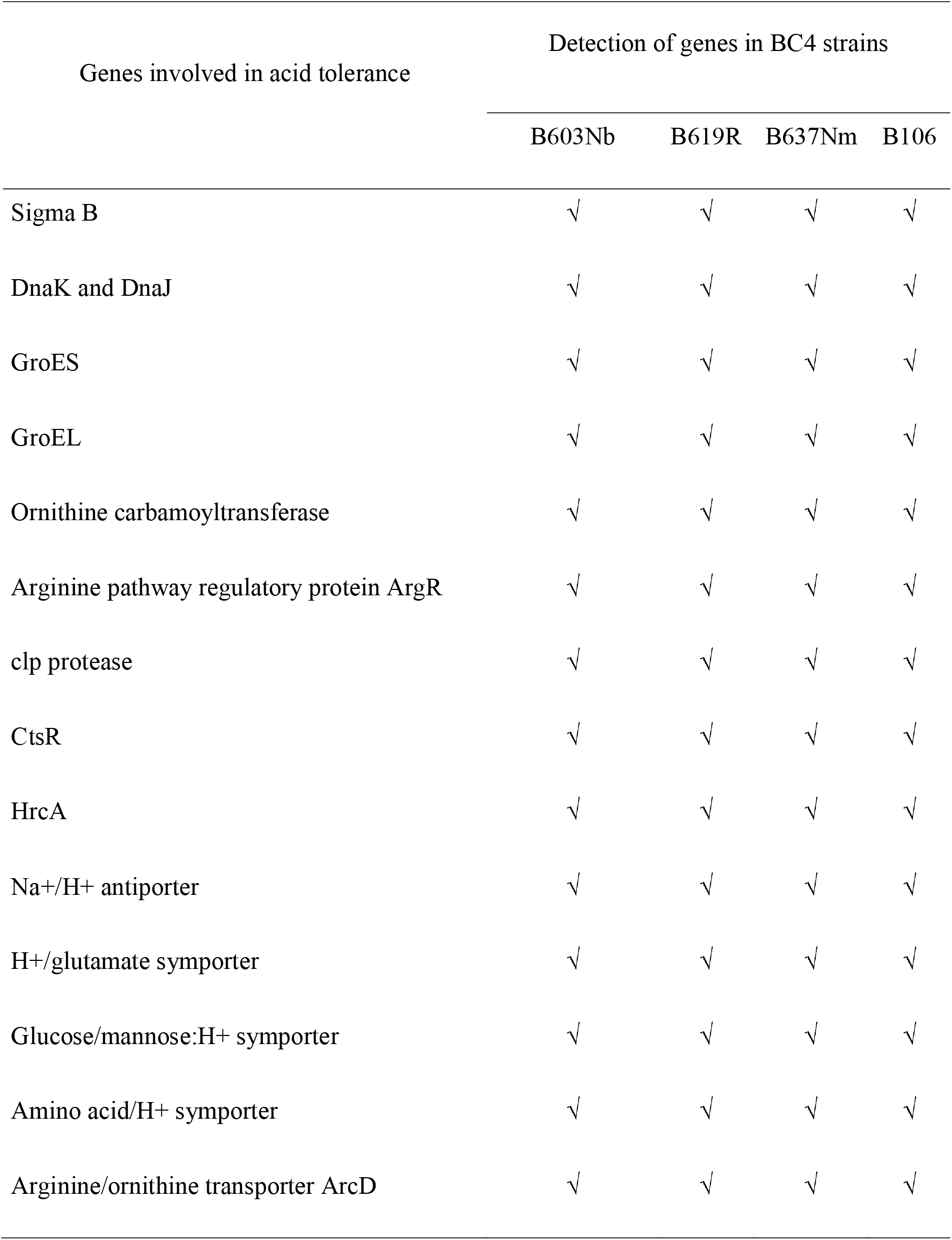
Detection of putative genes involved in acid tolerance in BC4 strains.

### Genes involved in tolerance to bile salt toxicity

Bile salts which are a result of cholesterol metabolism by hepatocytes, are excreted into the small intestine. They help in the formation of water-insoluble micelles with fatty acids originating from dietary fats. After their synthesis, they conjugate with either taurine or glycine, which makes them impermeable to the cell membrane and allows them to persist in the small intestine in higher concentrations. Bile salts also stimulate mucin secretion and bacteriostatic effects, which likely affect the intestinal flora (Hofmann, 1999). After encountering these conditions, the probiotic bacteria must reach the large intestine in significant numbers to allow further colonization and proliferation. The effects bile salts have on the intestinal microbiota can be detergent action, DNA damage, oxidative stress, and osmotic stress. Bile acids possess antimicrobial activity owing to their ability to act as detergents. If bile salts are present in high concentrations, they can rapidly dissolve membrane lipids and dissociate integral membrane proteins. Unconjugated bile acids can flip-flop across the cell membrane and enter the cell and cause damage to macromolecules. Bile salts are known to cause secondary structure formation of DNA. They also are involved in misfolding and denaturation of various proteins. Probiotics like various species of *Lactobacillus* have shown tolerance to bile with the help of proteins like bile salt hydrolase (BSH) and chologlycine hydrolase (Begley et al., 2005). However, both proteins were absent in the genomes of strains B603Nb, B619R, B637Nm, and B106. Reports have been published which say that the inactivation of genes that encode bile salt hydrolase (bshA and bshB) do not affect the bile tolerance property like in *L. acidophilus* and some strains of *L. plantarum* (McAuliffe et al., 2005). Proteins from the bile acid symporter family were detected, which regulate the uptake of bile. One major putative mechanism by which the four strains studied in the present investigation can show bile tolerance is sporulation. Sporulation has been known to provide a unique mechanism for bacteria to survive and exert their pathological effects in the intestinal tract (Yasugi et al., 2016). Seventy-five genes that are involved in sporulation were found in the genome of all four strains. Bacteria capable of sporulating often do so when they are faced with extreme conditions like starvation, extreme chemical environment, extreme temperature or pH, etc. This ability helps them survive in unfavorable conditions (Müller et al., 2014). The presence of these genes, especially the ones involved in sporulation, provides strong evidence to support the idea that the *A. clausii* strains B603Nb, B619R, B637Nm, and B106 can survive the bile toxicity encountered in the gastrointestinal tract of the host.

### Colonization of intestinal epithelium

One of the most important and essential properties of a probiotic strain is its ability to adhere to and colonize the intestine. Various health effects have been associated with the property of adhesion. Good adhesion to the intestinal mucosa is required to withstand the high flow rates observed in the small intestine. A major advantage of the adhesion of probiotics to the intestinal mucosa is that it can prevent the binding of pathogens, although no definite correlation seems to exist (Bibiloni et al., 1999). As opposed to that, if the probiotics have a higher affinity for the receptors present on the mucosa than pathogens or if they are simply present in a higher concentration than the pathogens, the probiotic bacteria may be able to displace the adhered pathogenic bacteria (Lee et al., 2002). The intestinal mucosa can be divided into three components, the epithelium, the lamina propria, and the muscularis mucosae. The epithelium is a single-cell layer lining the interior lumen of the gastrointestinal tract. Lamina propria is an interstitial tissue with a rich vascular and lymphatic network and abundant leukocytes. While the muscularis mucosae is composed of smooth muscle fibers. The interaction between these three compartments is dynamic in nature. The thickness of the mucus tends to vary from 120 µm to 830 µm. Due to the flow rate in the intestine (1m/h to 5-10 cm/h), it is important for adhesion for the probiotic bacteria to persist (Ouwehand & Salminen, 2003). After mining the genome of the *A. clausii* strains B603Nb, B619R, B637Nm, and B106, we found multiple genes encoding for proteins that help in the mechanism of adhesion and aggregation. The list of genes encoding for said proteins is summarized in Table 6. The presence of these genes indicates that the four strains studied in the present investigation show that they are likely to possess the ability to colonize the intestinal epithelium of the host.

**Table 6:**
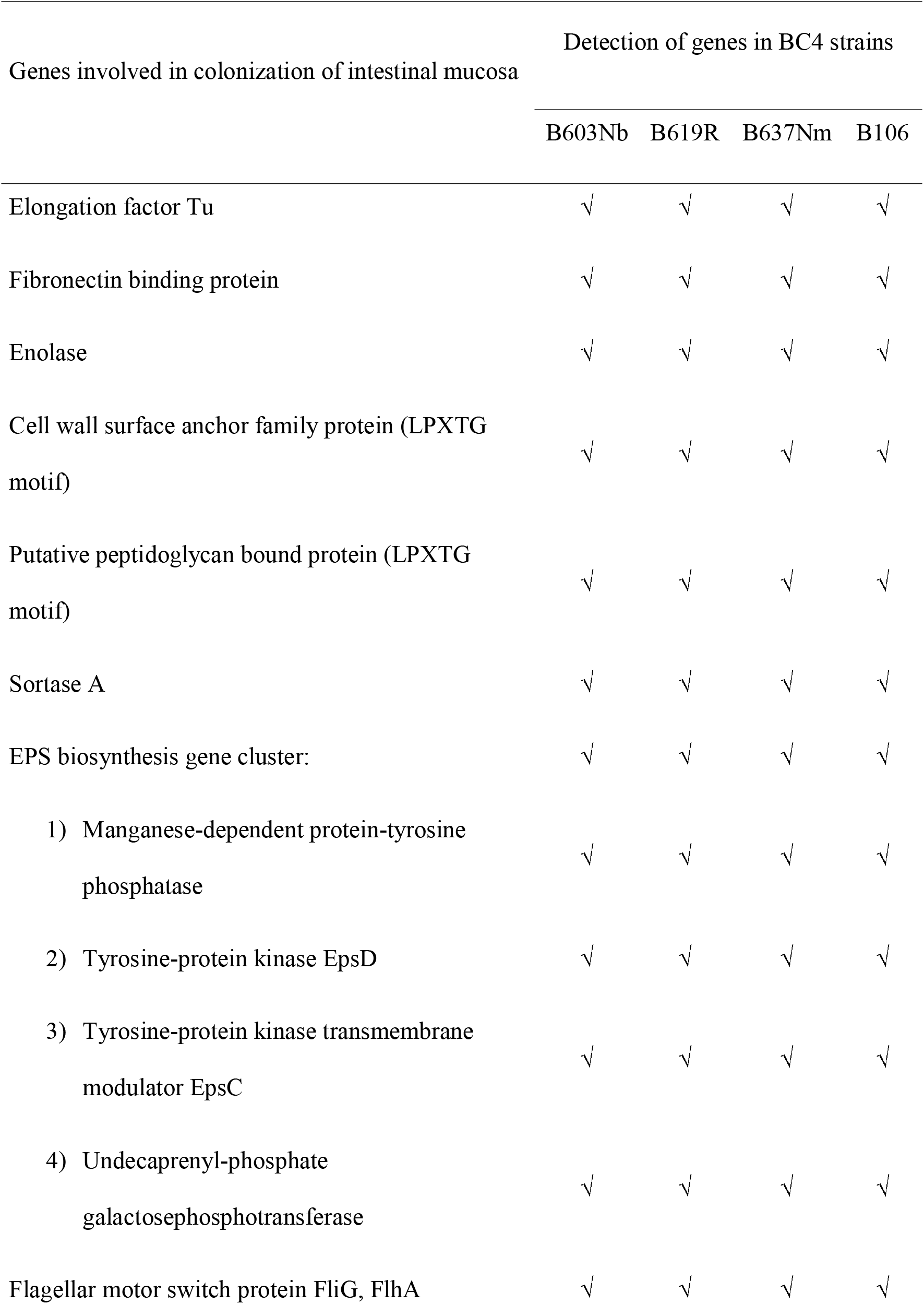
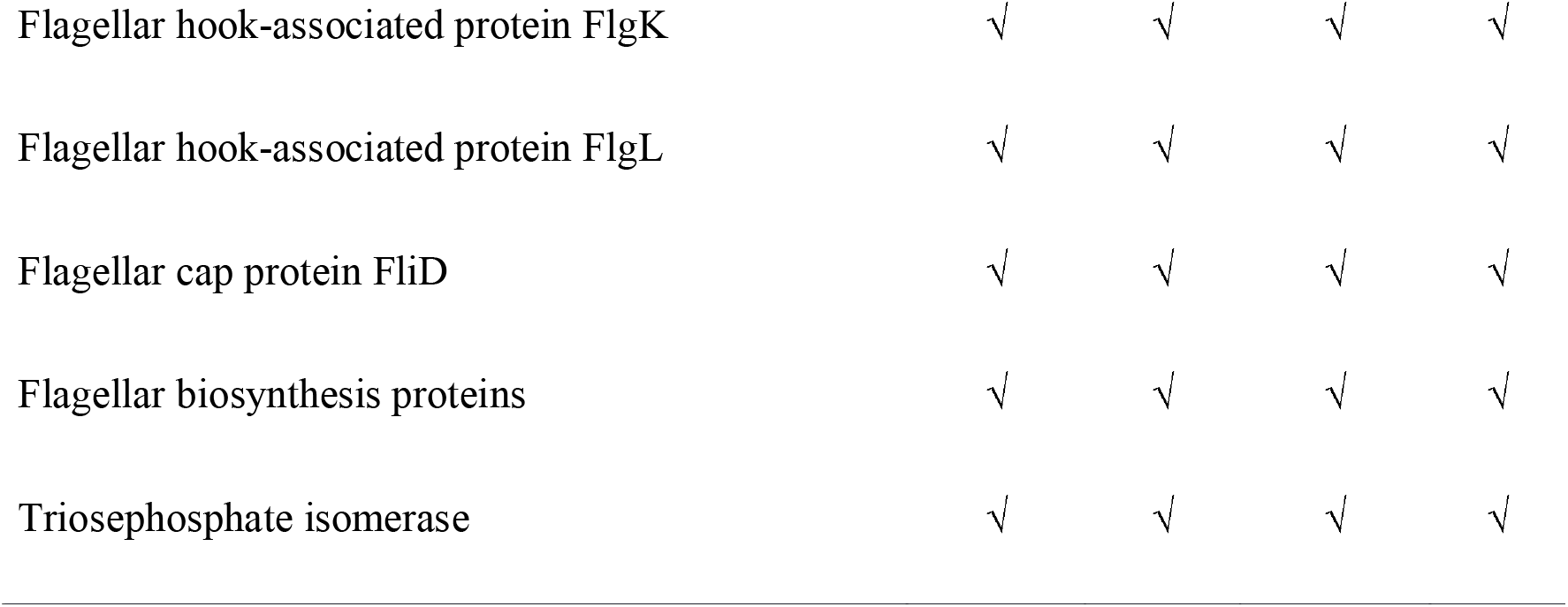
List of genes involved in colonization of intestinal mucosa.

The genes mentioned in the above table help with adhesion by various mechanisms. One mechanism employed by probiotics to adhere to host mucosa is the use of mucin/fibronectin-binding proteins. Protein EF-Tu has been known to mediate the attachment of bacteria to the intestinal wall. EF-Tu binds to fibronectin which is a part of the extracellular matrix. Generally, a cytoplasmic protein, the localization of this protein to the cell surface can be explained with the help of LPxTG peptidoglycan motifs. Thus, the presence of ER-Tu, as well as the LPxTG motif, can indicate the presence of a cell surface EF-Tu which is imperative for the binding of probiotics to the intestinal epithelium or mucins (Granato et al., 2004). Genome analysis of all four strains showed the presence of both EF-Tu as well as a single copy of fibronectin-binding protein, both of which could aid in the adhesion of bacteria to the host epithelium and fibronectin, respectively. Another protein, enolase, a key glycolytic enzyme, has been reported to have plasmin (ogen) binding property if expressed on the surface of certain bacteria (Pancholi & Fischetti, 1998). Three proteins-Cell wall surface anchor family protein (LPXTG motif), putative peptidoglycan bound protein (LPXTG motif), and sortase A (LPXTG specific) were found to be present in the genome of all four strains. All three proteins are LPXTG type anchors, which are known to be involved in cell surface localization and peptidoglycan interaction (Kapse et al., 2018). Many probiotic bacteria produce exopolysaccharides (EPS) which are exocellular biopolymers. EPS has been known to assist in a variety of functions like protection against desiccation, extreme pH and temperature, actions of antimicrobial compounds, attacks from a bacteriophage (Forde & Fitzgerald, 1999), and phagocytosis (Angelin and Kavitha, 2020). EPS, along with proteins, have been noted to play an important role in the transitory colonization of the intestinal mucosa along with antagonism of the enteropathogens (Kapse et al., 2018). The genomes of probiotic bacteria show ubiquity concerning eps gene clusters which could indicate that these bacteria can produce EPS in the intestinal environment, and a high enough concentration could be reached locally (Ruas-Madiedo et al., 2006). After mining the genome, we were able to find four genes, Manganese-dependent protein-tyrosine phosphatase, Tyrosine-protein kinase EpsD, Tyrosine-protein kinase transmembrane modulator EpsC, and Undecaprenyl-phosphate galactose phosphotransferase belonging to the EPS biosynthesis gene cluster. A flagellum can also participate in the adhesion of bacteria. Flagellar motility is an important prerequisite for colonization and persistence of probiotics as well as commensals. After initial attachment, bacteria often tend to aggregate to increase their chance of survival and further colonization. Aggregation can be either auto-aggregation, where bacteria belonging to the same species form aggregates, or co-aggregation, where genetically distinct species aggregate together with the help of various molecules. Aggregation is mediated by adhesins, flagella, various surface-associated molecules, lipoproteins, etc. Flagellar proteins like Flagellar motor switch protein FliG, FlhA, which are known to be involved in flagella-mediated auto-aggregation (Hoeflinger & Miller, 2017), were detected in the genomes of all four BC4 strains. The expression of flagellar structures and adhesive structures does not happen at the same time to prevent simultaneous motility and attachment. Various genes associated with flagellar adhesion like Flagellar hook-associated protein FlgK, Flagellar hook-associated protein FlgL, Flagellar cap protein FliD along with several flagellar biosynthesis proteins were found in all four strains. Triosephosphate isomerase, which is known to act as a cell surface adhesion molecule, was also detected.

### Oxidative stress tolerance

Probable genetic mechanisms adopted by BC4 strains to counter oxidative stress were investigated by mining the genes encoding enzymes that quench oxidative stress. Table 7 illustrates such genes detected in BC4 strains. When a cell is subjected to conditions where the prooxidant and antioxidant balance is disturbed, the cell enters a state known as oxidative stress. This disturbance in the balance often results in DNA hydroxylation, Protein denaturation, lipid peroxidation, apoptosis, etc. This may ultimately compromise the viability of the cells. Such conditions often arise when bacterial cells are introduced to harsh environments like those encountered in the gastrointestinal tract. On exposure to bile salts or low pH, the electron transfer chain is severely disturbed. This leads to the formation of reactive oxygen species (ROS) like superoxide anion radicals, hydroxyl radicals, and hydrogen peroxide. ROS are highly reactive in nature and can modify other oxygen species, DNA lipids, proteins, etc. one of the major sources of ROS is mitochondrial respiration. Mitochondria, peroxisomes, cytochrome p450, NADPH oxidase, and COX (cyclooxygenase) are some of the endogenous sources for the production of ROS. Over the course of evolution, bacteria have developed enzymatic defenses along with repair mechanisms to protect them from oxidative stress. Superoxide is one of the most abundantly found ROS. Their breakdown is catalyzed by the enzyme superoxide dismutase, which converts superoxides into hydrogen peroxide (H_2_O_2_) and water (Wang et al., 2017). Further, the decomposition of hydrogen peroxide is carried out by the enzyme catalase and glutathione peroxidase (Waris & Ahsan, 2006) in water (Wang et al., 2017). In addition to that, probiotic bacteria have been known to produce certain metabolites like folate and glutathione, which show antioxidant activity. Glutathione peroxidase, along with the non-enzymatic antioxidant GSH eliminates radicals like hydrogen peroxides, hydroxyl radicals, and peroxynitrite. The universal stress protein family is a superfamily of proteins that are produced in response to oxidative stress, among others (Kvint et al., 2003). Another mechanism employed the thioredoxin (Trx) system. This system includes NADPH, thioredoxin reductase, and thioredoxin. This system functions through its disulfide reductase activity regulating the protein dithiol/disulfide balance. Thiol-dependent peroxidases (peroxiredoxins) require electrons to remove reactive oxygen and nitrogen species with a fast reaction rate which is provided by the trx system (Lu & Holmgren, 2014). Besides, several chaperon proteins like DnaJ and DnaK are also involved in the general stress response along with GroES/EL, clp proteases, CstR, HrcA (Mols & Abee, 2011). Based on the above-mentioned mechanisms, the proteins present in the Enterogermina isolates have been summarised in Table 7.

**Table 7:** Probable genetic mechanisms adopted by BC4 strains to counter oxidative stress.

| Genes encoding protein involved in quenching of oxidative stress | Detection of genes in BC4 strains |  |  |  |
| --- | --- | --- | --- | --- |
|  | B603Nb | B619R | B637Nm | B106 |
| Superoxide dismutase | √ | √ | √ | √ |
| Catalase | √ | √ | √ | √ |
| Glutathione peroxidase | √ | √ | √ | √ |
| Universal stress protein family | √ | √ | √ | √ |
| Thioredoxin reductase | √ | √ | √ | √ |
| Peroxidases | √ | √ | √ | √ |
| DnaJ/DnaK | √ | √ | √ | √ |
| GroES/EL, clp proteases, CstR, HrcA | √ | √ | √ | √ |
| Peroxidase stress regulator | √ | √ | √ | √ |
| Nitrate and nitrite reductase | √ | √ | √ | √ |

### Sporulation ability of BC4 isolates

Sporulation is a mechanism possessed by some bacteria to produce nearly dormant structures called spores. In these structures, bacteria store their genetic material (DNA) when conditions become lethal and inhospitable. The spore-bearing ability of BC4 strains was investigated by mining the genes encoding proteins involved in the sporulation process (Table 8).

**Table 8:** Essential genes for sporulation present in the genome of *A. clausii* strains B603Nb, B619R, B637Nm, and B106

| Essential genes for sporulation | Detection of genes in BC4 strains |  |  |  |
| --- | --- | --- | --- | --- |
|  | B603Nb | B619R | B637Nm | B106 |
| Sporulation response regulator (Spo0A) | √ | √ | √ | √ |
| Stage II sporulation protein P | √ | √ | √ | √ |
| Stage II sporulation protein M (SpoIIM),<br>protein D (SpoIID), protein B, protein P | √ | √ | √ | √ |
| RNA polymerase sporulation specific sigma<br>factor SigF | √ | √ | √ | √ |
| RNA polymerase sporulation specific sigma<br>factor SigE | √ | √ | √ | √ |
| SpoIIGA, SpoIIE | √ | √ | √ | √ |
| a prespore engulfment protein, SpoIIQ | √ | √ | √ | √ |
| RNA polymerase sporulation specific sigma<br>factor SigH | √ | √ | √ | √ |
| RNA polymerase sporulation specific sigma<br>factor SigG | √ | √ | √ | √ |
| RNA polymerase sporulation specific sigma<br>factor SigK | √ | √ | √ | √ |
| Chromosome-anchoring protein RacA | √ | √ | √ | √ |
| Cell division initiation protein DivIVA | ✓ | ✓ | ✓ | ✓ |

Spores are multilayered structures that can be maintained for extended periods. They generally consist of the outer spore coat, cortex, and inner core. The outer coat has been known to be composed largely of proteins like keratin. It also contains diaminopimelic acid (DAP) or hexosamine. Phosphorous content also plays a role in the dense outer coating (Warth et al., 1963). These factors allow spores to survive in adverse conditions like heat, desiccation, radiation, chemical exposure, etc. furthermore; these spores can resuscitate into metabolically active cells again once the environmental conditions are favorable. Thus, sporulation can help probiotic bacteria overcome barriers presented by gastrointestinal stress like low pH, bile toxicity, the action of proteolytic enzymes, etc. While production of the probiotic formulations, manufacturing commonly involves the exposure of organisms to high temperatures, desiccation, freeze-drying, high pressures, etc. These conditions often lead to the death of microorganisms present in the formulation and, thus, a decrease in viability. Therefore, production costs often increase while dealing with probiotics containing non-spore-bearing bacteria. As opposed to this, if bacterial spores are used in the preparation of the formulation, the durability and sturdiness of spores allow them to withstand and survive all the unit processes making them a simpler and more economical choice. The genomes of *A. clausii* strains B603Nb, B619R, B637Nm, and B106 were mined for the genes involved in sporulation and 75 genes were detected. The presence of these genes indicates the strains studied in the present investigation possess the ability to sporulate and survive the acidic pH in the GIT and reach the gut unharmed. Some of the genes essential for sporulation were detected in BC4 strains. These genes are enlisted in Table 8. One of the most important players in sporulation is the sporulation response regulator (Spo0A). This transcription regulator initiates the differentiation of vegetative cells into heat-resistant spores through phosphorelay. This triggers the transcription of several SpoII genes, which are developmental regulators that are involved in asymmetric sporulation division, which gives rise to forespore (prespore) and mother cell. After division-specific gene expression, event characteristics of the two cell types are initiated. They are directed by RNA polymerase sigma factors, SigF in the forespore, and SigE in the mother cell (Piggot & Hilbert, 2004). Many such genes (SpoIIGA, SpoIIE) were present in the genomes of the *A. clausii* strains. Stage II sporulation proteins M, B, D, and P were found present. These proteins have a role in the internalization of the forespore through the breakdown of peptidoglycans between the two membranes at the septum, which separates the forespore from the mother cell. The proteins P, M, and D are complexed together where protein M serves as the membrane anchor and recruits protein P to the septum, which in turn recruits protein D (Mitchell et al., 2019). Proteins RacA, DivIVA, and FtsZ, which are known to be involved in morphogenesis and chromosome partitioning, were found in the genomes of *A. clausii* strains B603Nb, B619R, B637Nm, and B106. The protein RacA binds to the chromosome, while DivIVA is a polar division protein that acts as a bridge connecting the forespore and mother cell (Piggot & Hilbert, 2004). In addition to the above-mentioned proteins, several other proteins that are involved in the process of sporulation were detected in the genomes. The presence of these genes is further proof of the robustness of BC4 strains as a probiotic.

### Production of bacteriocins and antimicrobial compounds

Bacteriocins refer to a diverse group of peptides that are ribosomally synthesized by bacteria and archaea. These peptides show antimicrobial activity. The presence of bacteriocins is considered a highly desirable trait to be found in a probiotic organism. One of the major functions of bacteriocin is that it can behave as colonizing peptides. This property can help them in their introduction to the intestinal mucosa and/or establish dominance over existing pathogens. Apart from antimicrobial activity, bacteriocins can also function by inhibiting competing strains or pathogens. They also function as signaling peptides where they utilize the mechanism of quorum sensing to signal other microbial cells or immune cells of the host (Dobson et al., 2012). The first bacteriocin described was colicin produced by *Escherichia coli* (Cascales *et al*., 2007). However, currently, the bacteriocins most studied are those produced by lactic acid bacteria (LAB), followed by those produced by industrially important *Bacillus* species. Many other antimicrobial substances that were not well characterized are known as bacteriocin-like inhibitory substances (BLIS) (Abriouel et al., 2011). The main classification scheme for antimicrobial peptides of ribosomal synthesis currently available is that of the LAB bacteriocins. Abriouel et al., (2011) proposed a classification system for bacteriocins produced by organisms belonging to the genus *Bacillus*. It proposed a three-class classification system; *Class I* includes antimicrobial peptides that undergo different kinds of post-translational modifications, which includes peptides with modifications typical of lantibiotics & peptides, which include other unique modifications. *Class II* bacteriocins include small (0.77–10 kDa), ribosomally synthesized, non-modified and linear peptides, which are heat and pH stable. It includes pediocin-like peptides with a conserved YGNGVXC motif near the N-terminus, thuricin-like peptides with a conserved DWTXWSXL motif near the N-terminus, and includes other linear peptides, such as lichenin produced by *B. licheniformis*, or cereins 7A and 7B. *Class III* includes large proteins (>30 kDa) with phospholipase activity, such as megacins A-216 and A-19213 produced by *Bacillus megaterium* strains.

Putative bacteriocins were predicted using BAGEL 4 (van Heel et al., 2018). The organization of genes involved in bacteriocin production by BC4 strains is illustrated in Figure 3. In the three genomes, BAGEL predicted two putative peptides; gallidermin & sactipeptide. Gallidermin belongs to Class I lantibiotic, which specifically binds the cell wall precursor membrane-bound lipid II resulting in pore formation in the membrane and simultaneously inhibiting peptidoglycan biosynthesis. The 44 amino acid length query sequence of gallidermin showed 100% similarity with gallidermin/nisin family lantibiotics. The following amino acid sequence was obtained, which was similar to that of a query sequence obtained from the organism *Staphylococcus aureus* (strainMW2) MEKVLDLDVQVKANNNSNDSAGDERITSHSLCTPGCAKTGSFNSFCC. Nisin is a flexible molecule with antimicrobial activity against several Gram-positive bacteria like *Staphylococcus aureus, Listeria monocytogenes, Clostridium botulinum*, etc. (Kapse et al., 2018). Nisin displays pore-forming activity. Gallidermin is known to not only inhibit the growth of *Staphylococcus aureus* and *Staphylococcus epidermidis* but also prevent the formation of biofilm by both species (Saising et al., 2012).

Sactipeptides are a group of bacteriocins that have a characteristic thioether bridge (sactionine bond). This thioether bridge is installed post-translationally, and its presence is imperative for the antimicrobial activity of sactipeptides. The presence of these genes indicates the potential of *A. clausii* strains B603Nb, B619R, and B637Nm to possess antimicrobial activity, which can aid them in the colonization of the gastrointestinal tract and pathogen exclusion. It also provides insights into the putative mechanism by which they can fight the disease-causing pathogens like *Helicobacter pylori, Gardenerella vaginalis, Listeria monocytogenes, Clostridium difficile, Staphylococcus pneumonia*, and *Streptococcus aureus* (Cotter et al., 2013). In B106, BAGEL predicted two bacteriocins, class I Lantipeptide and the other from class Sactipeptide. The class I lantipeptide was identified as Gallidermin/ Nisin, a polycyclic antibacterial peptide commonly produced by the bacterium *Lactococcus lactis* (Kapse et al., 2019). However, to confirm the somewhat hypothetical evidence provided by genomic analysis, in vitro studies were carried out to see if *A. clausii* strains B603Nb, B619R, B637Nm, and B106 have the ability to inhibit the growth of pathogenic organisms. As all four strains show extremely high similarity both phenotypically and genotypically, strain B106 was selected as a representative of all four strains and was used to study the antimicrobial activity. Evaluation of antagonism towards pathogenic strains by agar overlay assay. Three sets were made to check the antagonistic activity against each pathogen. Each set had different incubation times i.e., 24 hours, 48 hours, and 72 hours. No inhibition was observed against the gram-negative pathogens which were tested. However, B106 showed to inhibit the growth of *Streptococcus aureus* after 72 hours of incubation. This preliminary screening provided further proof that *A. clausii* strains do have the ability to inhibit the growth of pathogens. Further study on the nature and characterization of the bacteriocin is pending.

### Production of hydrogen peroxide

The production of hydrogen peroxide (H_2_O_2_) is a known trait possessed by the bacteria residing in the gastrointestinal tract. The production of hydrogen peroxide may help probiotic bacteria establish dominance over the other residents of the GIT. H_2_O_2_ is a key player in causing oxidative damage to bacteria, especially those who do not possess hydrogen peroxide-scavenging enzymes like catalase and NADH peroxidase. There have been reports that the production of hydrogen peroxide is beneficial in more than the above-mentioned way. One report proposes that H_2_O_2_ has a homeostatic effect on the microbiota of the intestine, though the mechanism for this property is highly unknown (Hertzberger et al., 2014). Generally, the genes involved in hydrogen production are those encoding pyruvate oxidase, NADH oxidase, and lactate oxidase. The genomes of *A. clausii* strains B603/Nb, B619/R, B637/Nm, and B106 (Kapse et al., 2019) were mined to check the presence of genes involved in H_2_O_2_ production, and the following genes were found present. Even though the other essential oxidase genes were absent, the presence of NADH flavin reductase makes *A. clausii* strains a potential probiotic candidate with pathogen exclusion properties

## Conclusion

The present study in addition to the study carried out by Kapse et al. (2019) has reported insights into genome characteristics of four *A. clausii* strains constituting BC4 formulation. BC4 strains were unequivocally identified up to species level using a polyphasic identification approach. Each of the four BC4 strains was resistant to multiple antibiotics; however, the antibiotic resistance pattern of each strain was distinct from the other. The genome insights confirmed the ability of BC4 strains to resist multiple antibiotics. It was confirmed that the antibiotic resistance genes in BC4 strains were neither acquired nor transferable. The genome analysis did not reveal the presence of genes encoding any of the known emetic toxins. Further, genomes of none of the four strains possessed pathogenic islands or virulence factors, thus underscoring the safety of using each of the four isolates as probiotic oral supplements. Spores of all four strains were able to resist the harsh gastric transit. All four strains possessed adhesion molecules that helped in the colonization of the human gut epithelium. BC4 strains also possessed the ability to tolerate oxidative stress and produce bacteriocins. Thus, the extensive genetic characterization of the four BC4 strains underscored the potential of each isolate as a ’safe to use oral probiotics’ Differences in the antibiotic resistance profile also highlighted the fact that the combination of all four isolates is required to revive healthy gut microbiota when different combinations of antibiotics are used in the treatment.

